# Agnostic Tools for Life Detection: A Multiscale Approach to Biological and Chemical Reaction Networks

**DOI:** 10.1101/2025.08.13.670162

**Authors:** Bruno Cuevas Zuviría, Zach Adam, Cathryn Sephus, Daniel Cove, Betül Kaçar

## Abstract

Life manifests itself across a variety of different operative scales, ranging from molecular networks up to ecologies. Deep conceptual difficulties arise when trying to define exactly what life’s distinguishing attributes are, and how they may be quantitatively distinguished from abiotic systems that are complicated but demonstrably not alive. Here we assembled different biotic and abiotic chemical reaction networks to test whether network-level (macroscale) and reaction-level (microscale) metrics can impartially distinguish biotic from abiotic category networks. Macroscale attributes such as statistical tests for a discrete network’s heavy-tailed connectivity distribution generally distinguish *bona fide* biotic and protobiotic networks (*E. coli* heterotrophy, pruned protometabolism, KEGG) from abiotic networks but with two significant exceptions (radiolytic network and the Open Reaction Database). Microscale attributes such as an analysis of the frequency of a set of six primitive reaction motifs shows a depletion of some motifs in all chemical networks and the prevalence of others in specific networks; all networks were readily distinguishable from random network variants across all motifs. A comparison of the chemical spaces spanned by the different networks points to similarity and dissimilarity relationships between networks. Independent component analysis of an aggregate of all measured attributes, across all contexts, reliably distinguishes abiotic, prebiotic and biotic networks from one another. The combination of macro- and microscale attributes forms an agnostic toolset that may help to evaluate the prebiotic plausibility of different candidate settings for the origins of life, and may inform new ways of detecting and recognizing living systems (engineered or extraterrestrial) that differ from Terran biochemistry.

## Introduction

> “All happy families are alike; each unhappy family is unhappy in its own way.”

> - Tolstoy, *Anna Karenina*

We have only one systemic example of life on our planet, with all known forms descending from the Last Universal Common Ancestor (LUCA)^1,2^. Contemporary definitions of life have focused upon behavioral attributes (evolvability, variability, heritability, *etc*.; (Joyce, 1994)) rather than composition^3–5^. This approach affords some desired flexibility, but also introduces new problematic questions such as determining exactly what these specific attributes ought to be^6–12^. It is even less clear how these behavioral attributes, which are already difficult to quantify for readily observable biological examples, may be mapped to the likely possibility that other living systems beyond Earth that may deviate from Terran biochemistry.

It is common to refer to this problem as the “N=1 Problem”, the statistical difficulty of generalizing a universal phenomenon from a single sample. By comparison, there are countless variations of chemical systems at all scales (laboratory experiments, planetary geochemistry, atmospheric reservoirs, etc.) which may be *complicated* but are not *alive*. Each non-living system is arguably unique in its own ways of failing to be alive; a chemical recasting of the ‘Anna Karenina principle’ noted in other disciplines^13^ . This reframing presents a novel formulation of an inverted problem for non-living systems: the difficulty of generalizing a universal phenomenon from a nearly infinite number of contrapositive examples. Yet, this inversion can turn a large number of contrapositive examples from an onerous burden into a logical and statistical basis for comparison. An array of very different attributes found in living systems can be measured, and there is a high probability that a non-living system will be deficient in at least one or more of these detectable attributes. When both living and non-living systems are described and analyzed in a consistent manner, they may all be tested, compared and evaluated accordingly.

A robust contrapositive comparative analysis strategy would assist the assessment of settings and conditions in which life is thought to have started, but implementing such an effort carries its own logistical challenges. Where, amongst infinite possibilities, does one begin? Systems chemistry experiments consistent with our geological understanding of the early Earth have uncovered plausible pathways leading to many of the molecules found in living organisms, including intermediate metabolites^14–18^, compartmentalization^19–22^, and the components of information polymers^23–29^. The synthesis of many important biomolecules has been attained by synthetic procedures that emulate prebiotic chemical reactions, but there is not yet a full developed theory or experiment that shows how to turn these biomolecules into self-organizing and evolving entities with life-like behaviors ^30,31^. Developing the logical formulation above into a practical research question demands a sweeping and consistent description of many different chemical systems, in a variety of operative contexts (most of which are not associated with life), and at a variety of different scales to serve as a statistical testbed for evaluating hypotheses regarding detectable attributes of life.

Here, we compile 19 chemical reaction networks (CRNs) describing a variety of very different geochemical (common redox reactions for Earth minerals, hydrothermal vents), atmospheric (Titan, exoplanetary hot Jupiter, solar system gas giants, Martian surface chlorides), cosmochemical (protoplanetary disc, with and without surface chemistry), biotic (*E. coli* metabolism, *E. coli* translation, *Thermotoga maritima* metabolism, *Methanosarcina bakeri* metabolism, KEGG), protometabolic (ancestrally-inferred protometabolic, phosphate-free inferred protometabolic), prebiotic (surface radiolytic, formose reaction, and aggregated prebiotic chemical reactions) and chemical engineering (combustion processes, Open Reaction Database) contexts. All network information is tabulated in the Supplementary Information and through an interactive site (https://chemorigins.bact.wisc.edu/crna). For each contextual setting, we have extracted the chemical reactions that they describe, homogenizing each network’s information into computer-readable formats. The resulting dataset of chemical reactions was analyzed using different analysis techniques to compare candidate attributes that distinguish biotic from abiotic contexts.

The network analyses evaluate attributes at both the macroscale (network topology such as connectivity distribution scale factor and mass modality) and microscale (intra-reaction patterns between compounds that form primitive reaction motifs). The metrics studied here (heavy-tailed distribution, mass modality/discontinuity, reaction micro-motifs, and chemical identity) form a basic toolset that usefully compares and contrasts biotic and abiotic chemical reaction networks.

### Networks

We consider a CRN as a directed bipartite graph containing two types of nodes: reactions and molecules. Each reaction must be connected with molecules through incoming edges (reactants) and through outgoing edges (products). This simplified description has proven useful to investigate the evolutionary mechanisms behind the emergence of metabolism^32^, the topological signatures of chemical systems^33,34^, and the principles shaping the design of organic synthesis pathways ^35^. Additional details such as rate constants and reaction conditions enable complicated simulations of systems chemistry and dynamics, though this is beyond the scope of this effort.

Our focus is on real-world CRNs, and we describe the reactants and products of the different models using chemical identifiers to i) be able to combine information coming from different chemical reaction networks and ii) assess the molecular properties of each molecule. We attempted to obtain an InChI or SMILES description of each molecule’s ground state, except in those reactions that depend entirely on specific charged structures (*e.g*. ionization reactions). We obtained these computer-readable chemical representations by consulting different databases, cross-referencing with PubChem or KEGG, or, in some cases, through manual annotation. Some specific compounds are difficult to translate into these representations due to specific features (*e.g*. the notation of different electronic states) and some networks are not fully comparable in chemical terms (*e.g*., some molecules are macro-molecular complexes). **Table 1** summarizes all of the real-world CRNs that were assembled, compiled and analyzed in this study, and provides an overview of their key sources and attributes. The column in the right indicates whether the network chemical reactions were readable in SMILES format, and if not, why it could not be formatted.

**Table 1.**
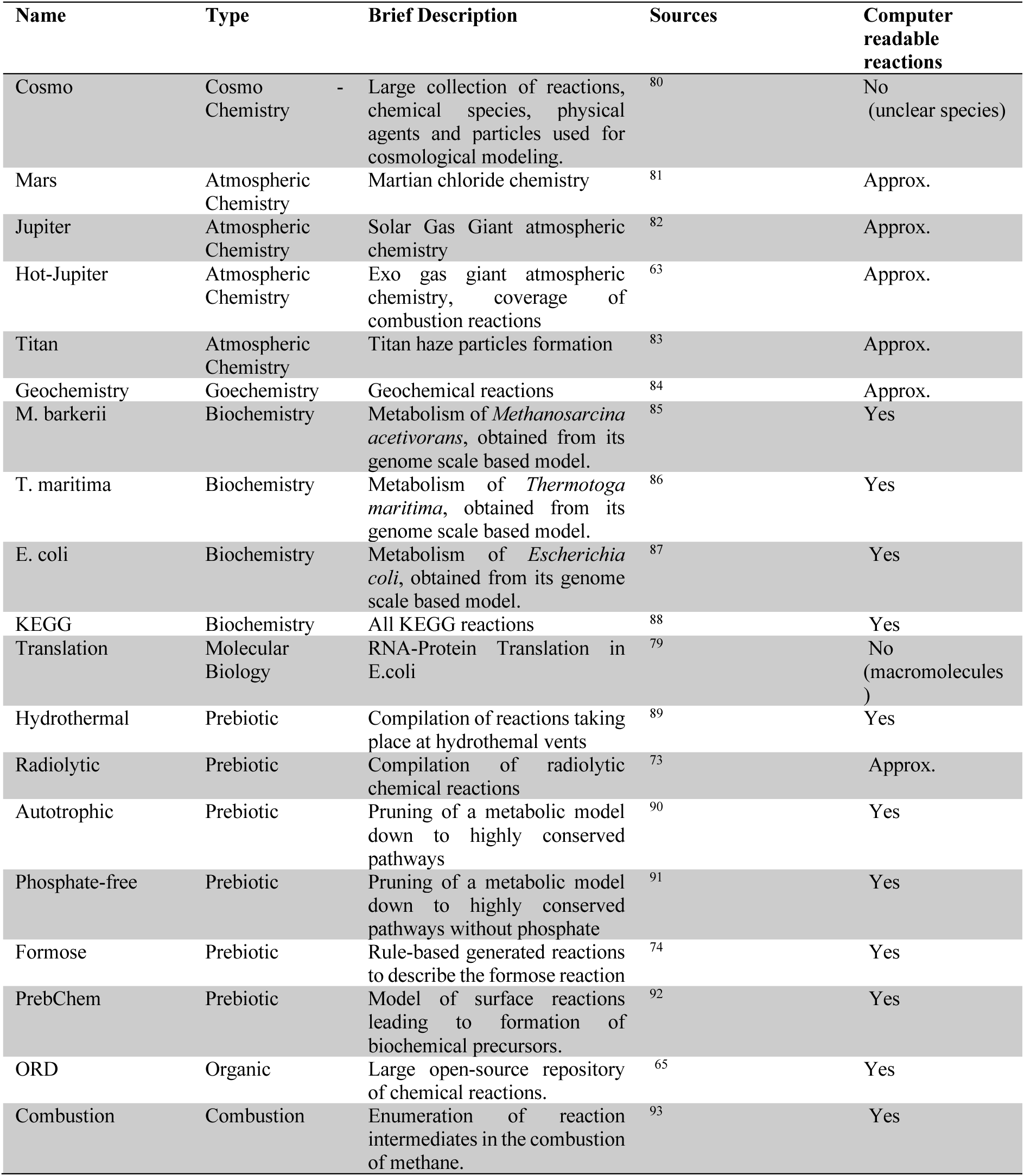
A tabulated summary of all chemical networks analyzed in this study. The right-end column indicates whether it was possible to systematically convert all species into computer-readable formats.

### Network analyses

Each set of chemical reactions generates a bipartite network. Previous analyses have shown how this kind of network has specific properties relevant to complex systems^34,36^. We characterize these networks at the macroscopic level using their degree distribution and mass-degree correlations and at the microscopic level of individual reactions using their reaction motif densities. The largest networks in our collection (the Open Reaction Database and the Cosmochemistry networks) were too large to conduct the micro-motif and chemical and reaction overlap analyses and were therefore omitted from the dimensional reduction Independent Component Analysis (see Multidimensional analysis section).

### Degree distribution

We define the degree of a node as the number of edges that start or end on a given node. This property has been studied in detail in previous work from the last two decades^32–34,36–44^ . Most chemical reactions involve only a few reactants and products, while molecules could potentially participate in many reactions. Therefore, we focus our analysis on the degree of molecules, which measures the number of reactions in which a molecule is produced or consumed. For each CRN, we use the degree value of all its compounds to generate a complementary cumulative distribution function (CCDF) to assess whether the network CCDF fit is more likely to match with a system with a homogeneous (i.e., exponentially-truncated) or heterogeneous (i.e., heavy-tailed) degree distribution. This analysis is carried out by fitting two different statistical distributions to the CCDF histograms: i) an exponential distribution [Eq1] which is more typical in homogeneous networks, and ii) a power law distribution, [Eq2], which is more typical in heterogeneous networks. Networks describing real-world systems can span a range of best-fits on a continuum from heterogeneous to homogeneous distributions, though, and a higher value for a degree distribution exponent can indicate that the probability of finding a highly connected node decreases rapidly (leading to a lower probability of the network being heavy-tailed)^45,46^ . We use the Python powerlaw package to infer which is the most likely fit for a network by comparing the log-likelihood ratios between distributions ^47^ . For each network, regardless of its likeliest degree distribution, we additionally calculate and plot the exponent α of the best fit of the CDDF to a power law distribution to test a hypothesis that the numerical value for a network’s degree distribution can sort between biotic and abiotic networks.

### Mass-degree correlation

CRNs involve chemical entities of different sizes, from elemental ions to large information polymers. While a complete description of all the objects and their internal states that make a given real-world system is intractable (e.g. a complex molecule with multiple folds, a hydrothermal vent, or a planetary atmosphere), we can consider a small number of objective attributes such as the mass of each of the molecules described in our CRNs. We computed the mass of those molecules using the rdkit package based on their molecular description. Note that some of the networks have molecules whose notation obfuscates their translation to computer-readable formats (e.g. SMILES, InChI), so we could not apply this specific analysis to all networks. The column of **Table 1** labeled ‘Computer Readable Reactions’ indicates those networks that had easily convertible formats, those which did not, and those for which as many molecules as possible were manually annotated (*approx* label). We correlated mass and degree using a simple linear regression on a log-log chart between masses and degrees of compounds. We test the hypothesis that the log-linear slopes of living systems have correlation coefficients between these variables that tend to cluster around characteristic values, while non-living systems do not.

### Micro-motifs

Micro-motifs are a way of counting the basic ways that any two or more objects in a network may be connected to one another. In this work context, two molecules may be connected by reactions to form a simple cycle between them, or part of a linear chain of compounds, or with parallel connections between their reactions. The total count of each connection type may be normalized by the total number of the network’s connections to measure the statistical distribution of how individual nodes are connected within a given network. Networks describing living systems may be more or less likely to contain a certain percentage of motifs compared to either non-living or randomized networks^48–53^ . Modern techniques enable the efficient counting of micro-motifs in small and medium networks. We focused on the connections that can be established in a network with 4 nodes: two molecules (A and B) and two reactions (r1, r2). Given that our CRNs are directional, we consider different types of motifs depending on the direction of the edges between the molecules and the reactions. We counted the number of motifs using DotMotif^54^. We also compared the results against randomized networks obtained by shuffling edges connecting reactants and reactions or reactions and products. This randomization ensures that the resulting random networks have the same size and degree as a given network, resulting in a controlled basis for comparison ^33^. We test the hypothesis that more structured micro-motif relationships (such as those that can lead to linear pathways or cycling between compounds found in biological networks) are statistically unlikely in random networks, and less structured relationships (such as refractory connections that do not create pathways or cycles) are more likely in random networks. We also test the hypothesis that an aggregation of multiple micro-motifs can usefully distinguish between biotic and abiotic networks through a multi-variate analysis (see *Multidimensional Analysis*).

### Chemical features

We quantified how chemical reaction networks span similar regions of chemical space (an abstraction to indicate the whole collection of possible molecules and reactions). We use two notions of overlap: hard, to quantify the overlap as the number of identical objects (reactions or molecules) shared between networks; and soft, to quantify the overlap as the number of objects that share some similarity. The computation of hard overlap took place by hashing molecules and reactions (mapping a molecule or a reaction to a code, so two molecules or reactions are unlikely to share the same code unless they are precisely the same). We use the InChiKey of each compound and its number of atoms to generate two hash keys depending on the size of the molecule. We disregard the protonation and charge states of the larger molecules by removing the last character of the InChiKey, while we conserve those for the smallest molecules (less than 6 atoms other than hydrogen). For reactions, we sort reactants and products and then apply a standard hashing protocol.

To study the soft overlap, we need to convert each reaction into a vector (embedding) that can be used to compute similarity relationships. We use the Transformers-based Embedding model RXNFP^55–60^, a neural network that converts a reaction SMILES sequence into a fixed-length vector that enables comparisons. To display the soft overlap, we employed a 2-D-based representation using dimensional reduction techniques known as t-distributed Stochastic Neighbor Embedding (t-SNE^61^).

### Multidimensional analysis

Each of the individual analyses conducted above provides numerical and potentially orthogonal dimensions that can be used for comparing and contrasting complicated CRNs. A multidimensional analysis technique can impartially parse these different data to evaluate which dimensional data (if any) distinguish the networks into insightful categories. In our case, the analysis can determine whether abiotic and biotic networks cluster together or apart from one another when both micro- and macroscale data describing their CRNs are processed. We use Independent Component Analysis (ICA) to reduce the dimensionality of our problem into a smaller manifold of potentially meaningful dimensions. ICA produces a set of vectors that statistically capture trends observed in the data. It differs from Principal Component Analysis (PCA), in which trend capture is not a product of a set of vectors but a rotation of the vector space. After normalizing the data, we employed the FastICA routine provided in ScikitLearn^62^. We test the hypothesis that an ICA informed by a multidimensional combination of the micro- and macroscopic CRN feature measurements describe above can reliably distinguish between biotic and abiotic networks.

### Interactive server

An interactive web portal based on the Python library *Streamlit* was constructed to enable users to both explore and download all of our source data. The portal address is https://chemorigins.bact.wisc.edu/crna. We also provide our pipeline code to enable users to perform similar computations.

## Results

### A new repository of chemical reaction networks

We were able to compile 19 distinct CRNs, providing the description of 167,889 reactions involving 155,736 compounds. Each network provides an incomplete snapshot of the chemistry that is possible within the system it describes. The networks also have different sizes, as shown in **Table 2**: from the enormous collection of reactions of ORD (100k in this paper; 2M reactions in the original network) to 80 in the smallest network we collected (hydrothermal network). To make this collection of networks available for future research, we have built an open repository of chemical reaction networks related with origins of life, which can be accessed at https://so23-chemical-reaction-networks.streamlit.app/.

**Table 2:** Network statistics: number of reactions, molecules, edges, and weak components (defined as components found within disconnected subgraphs within the main graph).

| Name | No. of Reactions | No. of Molecules | No. of Edges | No. of Weak Components |
| --- | --- | --- | --- | --- |
| Combustion | 434 | 33 | 1671 | 1 |
| ORD | 100000 | 143730 | 514481 | 348 |
| Cosmo | 48699 | 982 | 204576 | 1 |
| Mars | 277 | 51 | 1177 | 1 |
| Titan | 809 | 87 | 3062 | 1 |
| Jupiter | 963 | 97 | 3707 | 1 |
| Hot-Jupiter | 916 | 96 | 3526 | 1 |
| geochemistry | 204 | 29 | 1165 | 1 |
| hydrothermal | 80 | 86 | 295 | 1 |
| prebchem | 311 | 234 | 1015 | 2 |
| radiolytic | 541 | 220 | 2175 | 1 |
| formose | 107 | 50 | 391 | 1 |
| autotrophic | 377 | 337 | 1643 | 1 |
| phosphate-free | 315 | 219 | 1080 | 1 |
| T. maritima | 498 | 497 | 2302 | 13 |
| M. barkeri | 531 | 554 | 2533 | 22 |
| E. coli | 1583 | 959 | 6720 | 3 |
| KEGG | 9811 | 6997 | 42881 | 16 |
| Translation | 1433 | 478 | 8931 | 1 |

**Table 3.** A breakdown of network topology values for all considered networks. Alpha records the calculated power law exponent. <K> records the average connection of the network. P is the calculated Fischer’s significance value indicating a power-law fit (as a proxy for a heterogeneously connected network). P<0.05 indicates those that meet the significance threshold for testing for a heterogeneous distribution. Distribution records the determined likeliest fit for the raw network connection data.

| Name | alpha | $\langle k \rangle$ | p | p < 0.05 | Distribution |
| --- | --- | --- | --- | --- | --- |
| combustion | 5.49 | 50.64 | 3.85E-01 |  | other |
| ord | 2.15 | 3.58 | 1.07E-34 | * | power-law |
| cosmochemistry | 2.44 | 208.33 | 2.30E-03 | * | power-law |
| Mars | 3.44 | 23.08 | 2.73E-01 |  | other |
| Titan | 3.66 | 35.20 | 1.06E-01 |  | other |
| Jupiter | 3.62 | 38.22 | 5.10E-01 |  | other |
| hot-Jupiter | 3.40 | 36.73 | 8.52E-01 |  | other |
| geochemistry | 3.89 | 40.17 | 9.04E-02 | * | other |
| formose | 2.86 | 7.82 | 2.90E-02 | * | other |
| hydrothermal | 2.45 | 3.43 | 5.59E-3 | * | power-law |
| prebchem | 2.71 | 4.34 | 1.07E-01 |  | other |
| radiolytic | 1.85 | 9.89 | 4.43E-17 | * | power-law |
| autotrophic | 3.18 | 4.88 | 2.43E-16 | * | power-law |
| phosphate-free | 2.55 | 4.93 | 1.48E-02 | * | other |
| T. maritima | 2.43 | 4.63 | 1.46E-07 | * | power-law |
| M. barkeri | 2.48 | 4.57 | 3.43E-08 | * | power-law |
| E. coli | 2.33 | 7.01 | 4.48E-05 | * | power-law |
| KEGG | 2.37 | 6.13 | 1.17E-16 | * | power-law |
| translation | 3.53 | 18.68 | 3.25E-14 | * | power-law |

### Heavy-tail degree distributions distinguish complex chemical reaction networks; power-law exponent can cluster by network category

There is a large and nuanced body of work exploring the principles that give rise to chemical networks characterized by heterogeneous degree distributions — networks where most nodes have very few connections, and a small set of nodes have an enormous number of connections^32^ . We analyzed the degree distributions across the different chemical reaction networks, obtaining two kinds of behaviors: networks where the degree distribution is more likely to be a homogeneous degree distribution and networks more likely to fit with a heterogeneous distribution. The adequacy of either heterogeneous degree distributions or to homogeneous degree distributions can be numerically estimated using log-likelihood ratio tests to calculate the p-value ^47^ . Plots that show the Complementary Cumulative Distribution Function (CCDF) versus bins of connectivity (k) are useful for visually comparing homogeneous and heterogeneous fits to the network data.

For example, the CRN describing a Hot-Jupiter exoplanet model^63,64^ , can be reasonably fit by either homogeneous or heterogeneous distribution types (**Figure 1a**) and the test for a heterogeneous fit is inconclusive. This contrasts with the CRN describing our subset of the Open Reaction Database network ^65^ , which shows a much better fit of a power-law degree distribution (red line) than for a fit of an exponential model representing the degree distribution of a homogeneous degree distribution (dotted grey line) (**Figure 1b**), therefore a heterogeneous fit is more likely. This analysis was conducted for all networks, and the results for each network’s average connectivity (<K>) were plotted versus each network’s best-fit connectivity distribution slope (𝛂) (**Figure 1c**). The marker for each plotted network was colored according to the network category; was sized according to the overall size of the network (as measured by the number of unique chemical compounds it contained); and the shape of each marker indicates whether their respective network power-law versus exponential log-ratio test was significant (power-law most likely fit and with p<0.05 are marked by an X, all others are marked O).

**Figure 1:**
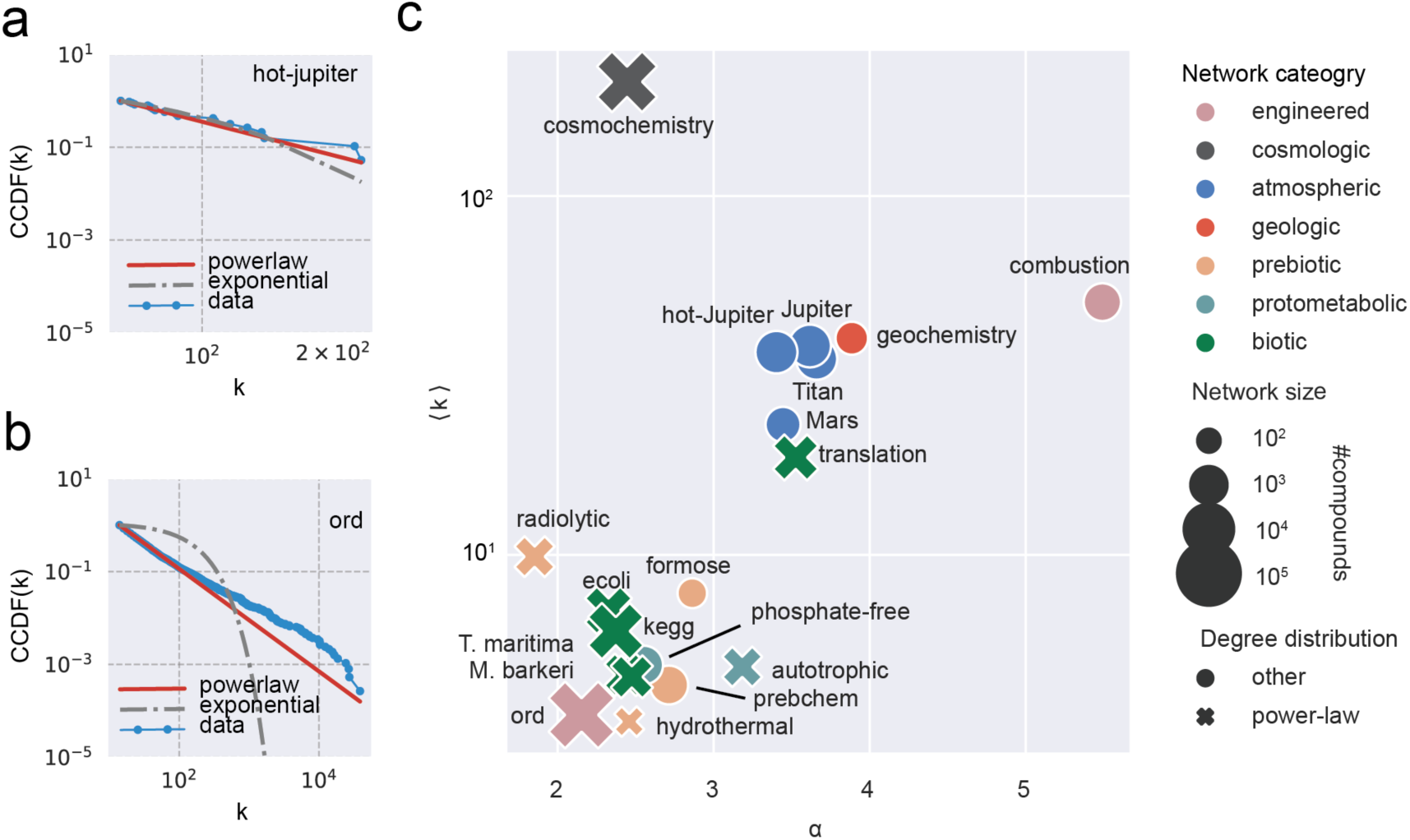
Heterogeneous degree distributions. **a)** The Hot Jupiter chemical reaction network’s degree distributions are represented by its complementary cumulative density function (CCDF) versus degree k. Homogeneous and heterogeneous fits corresponding to exponential and power-law degree distributions, respectively, work equally well. **b)** Same as a), but for the ORD chemical reaction network. A power-law degree distribution is a better fit. **c)** Power-law fit exponent 𝛂 versus average degree. Marker sizes describe the size of the network and marker shapes describe the most likely distribution.

In the resulting plot, there is a clear difference between networks containing a large number of gaseous compounds (atmospheric and cosmochemical) and the other chemical networks. Gaseous networks have high average connectivity degree <K> and high power-law exponent fit 𝛂, while all other networks generally show much lower average degrees and lower power-law exponents. The Cosmochemistry network was also the only such network to have a likely heterogeneous distribution.

### Biotic networks have low correlations between log-mass and log-degree variables

Due to thermodynamic laws, every active chemical system takes energy (e.g. light, hydrogen, fuel) from the environment and returns it to the environment as degraded quality energy (e.g. infrared radiation, heat, CO2)^66^ . Quantifying how this transformation occurs could be a key aspect differentiating biotic and abiotic chemical systems. Life consumes energy to generate chemical systems with a higher degree of organization and specificity, while most abiotic processes have been characterized as the opposite ^67^ . Knowing the specific energy, abundance, and flux of each chemical would lead to a fuller understanding of the energy transformations through a chemical system. Unfortunately, near-complete descriptions of these kinds of data are only available for relatively simple systems.

To bypass these limitations, we assume two very basic principles to analyze the given networks: (1) larger molecules (particularly organic molecules) generally have higher standard molar enthalpy of formation values and therefore require more energy to produce from substrates^68^ , and (2) the number of reactions involving a given compound is correlated with its time-averaged abundance in that system. Real individual molecules and chemical systems often deviate from these assumptions in different ways, but the assumptions are simple enough that they permit a basic assessment of energy distribution in a system without reference to the idiosyncrasies of its constituent molecules or the rate constants of their reactions within the network.

In **Figure 2** the log-degree of each compound in a network is plotted against its log-mass. From these plots we calculate a best-fit linear relationship between these values, as well as a Pearson’s correlation coefficient (r) analysis to quantify the correlations between these two variables. The shape of the distributions shows important differences between the different kinds of networks: nearly all biotic networks have r values near 0 (indicating very little correlation), while most systems that involve radicals (atmospheric networks and the radiolytic prebiotic chemical network) show a moderate negative correlation, with values between -0.5 and -1 (**Table 4**). In the largest chemical networks, most of the molecules cluster around a 100-500 Da region with very few connections, which could correspond to secondary metabolites or end products in organic chemistry.

**Figure 2.**
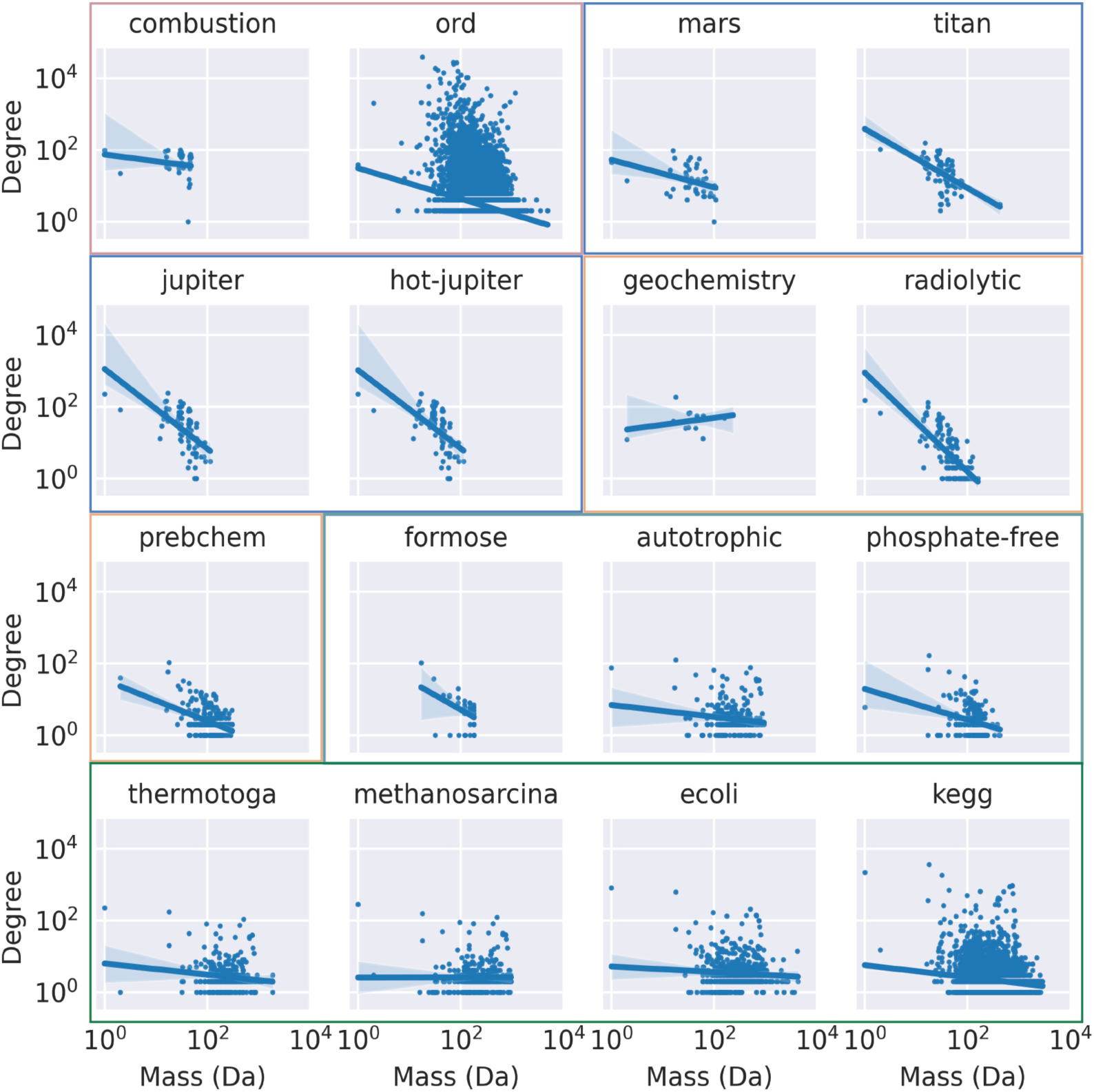
Molecular degree-mass distribution across the different networks analyzed. The line represents a least-squares linear fit between the log-mass and the log-degree distributions.

**Table 4.** Regression analysis of log-mass/log-degree relationship. R denotes the Pearson’s correlation coefficient. Slope denotes the calculated slope of the least-squares linear fit between these two variables for each network. The p-value represents the calculated Fischer’s significance test. *Translation* and *Cosmochemistry* networks were not included here as the reactions were not computer-readable (see Table 1).

| network | r | slope | P value |  |
| --- | --- | --- | --- | --- |
| **combustion** | -0.17656 | -0.1884 | 3.26E-01 | p > 0.01 |
| **ord** | -0.24949 | -0.3439 | 0.00E+00 | p < 0.01 |
| **Mars** | -0.36850 | -0.3845 | 1.78E-02 | p > 0.01 |
| **Titan** | -0.58601 | -0.8248 | 2.68E-08 | p < 0.01 |
| **Jupiter** | -0.59628 | -1.1124 | 2.72E-09 | p < 0.01 |
| **hot-Jupiter** | -0.59425 | -1.0905 | 7.76E-09 | p < 0.05 |
| **geochemistry** | 0.29791 | 0.1914 | 3.23E-01 | p > 0.05 |
| **formose** | -0.43745 | -0.8114 | 1.49E-03 | p < 0.05 |
| **hydrothermal** | -0.435260 | -0.5099 | 2.81E-5 | p < 0.05 |
| **radiolytic** | -0.71318 | -1.3810 | 1.64E-23 | p < 0.05 |
| **prebchem** | -0.40318 | -0.5718 | 1.47E-10 | p < 0.05 |
| **autotrophic** | -0.18636 | -0.1876 | 5.85E-04 | p < 0.05 |
| **phosphate-free** | -0.28842 | -0.4277 | 1.45E-05 | p < 0.05 |
| **T. maritima** | -0.13973 | -0.1541 | 8.66E-03 | p < 0.05 |
| **M. barkeri** | 0.00205 | 0.0021 | 9.68E-01 | p > 0.05 |
| E. coli | -0.07418 | -0.0785 | 5.50E-02 | p > 0.05 |
| KEGG | -0.14179 | -0.1682 | 1.09E-31 | p < 0.05 |

### Biotic networks tend to have more parallel reaction motifs and fewer refractory motifs than abiotic ones

Network motifs are elementary interaction modes between nodes that underpin important features of dynamical systems, such as feedback loops or cycles. We have considered all the possible motifs that can be built by two reactions related to two different molecules (**Figure 3**), and we have searched for them in our networks using the search method implemented in DotMotif^54^ . The first type (*m0*) relates to small minimal cycles—which, in some cases, could be autocatalytic. Motifs *m1* to *m4* describe reactions or molecules that interact to form symmetric patterns: serial reactions, parallel reactions, shared products and shared reactants. The *m5* and *m6* represent refractory interactions that lack symmetry or overall structure, and we aggregate the counts for both under the label *mx*. Our objectives are to compare and contrast living and non-living system networks to each other, and also to compare both of these kinds of natural networks to randomized networks.

**Figure 3:**
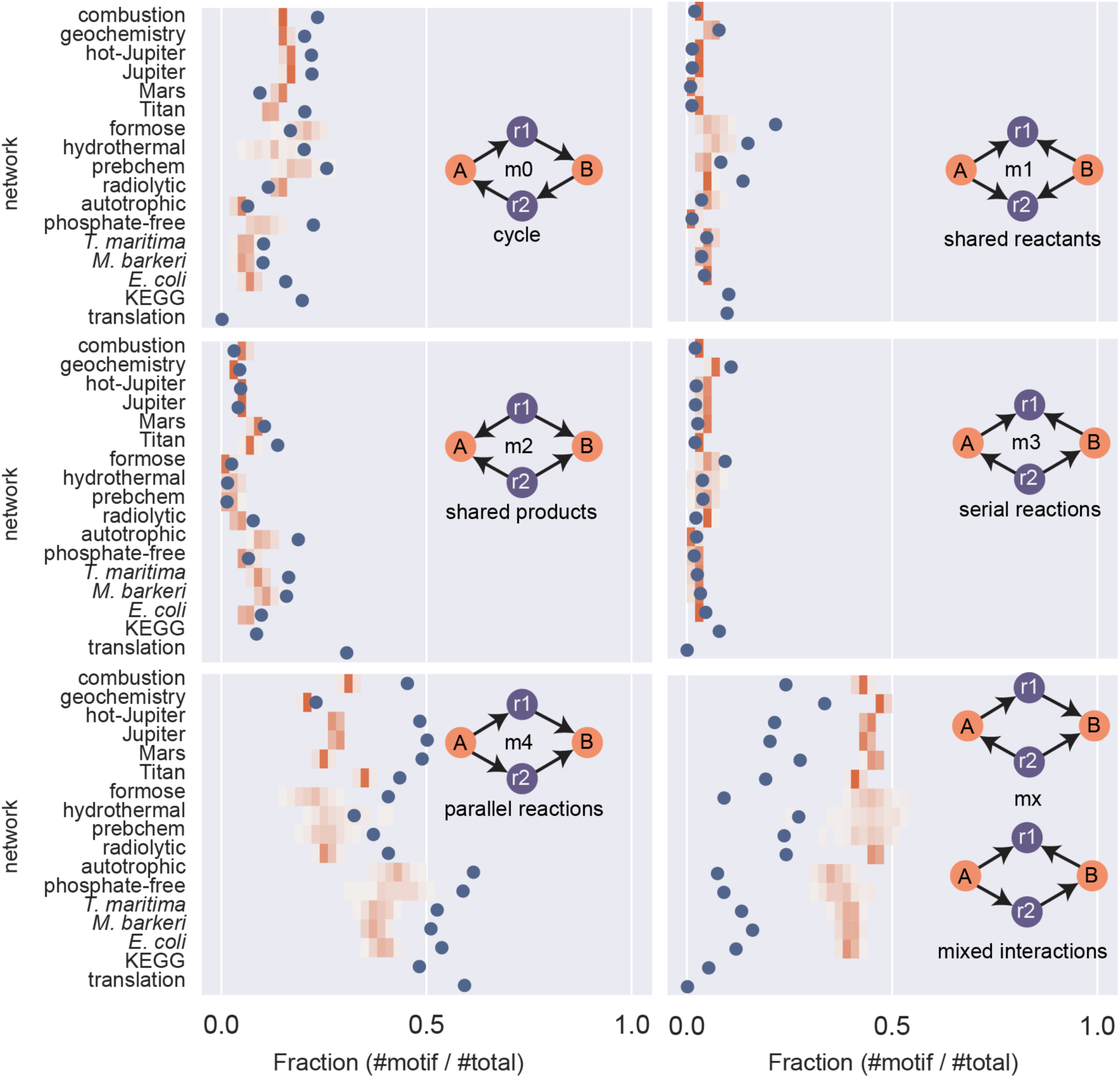
Frequency of each possible two-compound, two-reaction motif in each named bipartite graph. Blue dots denote the quantified frequency of each motif within the canonical network, while the orange shade indicates the frequency and distribution of the motif in randomized variants of the same named network. Note that the KEGG and translation networks were too large to apply, analyze and quantify randomization effects, and the ORD and cosmochemistry networks were too large to definitively measure motif frequencies and randomization effects.

In **Figure 3**, we compare the fraction of each motif in each network against those found in 40 randomizations of the same network. The fractions of most micro-motifs found in natural networks are similar to their randomized counterparts for motifs m1, m2 and m3, but there are striking differences between natural and randomized networks for motifs m0, m4 and mx. Natural system networks (biotic and abiotic) tend to contain more cyclic and linear connections, while they contain far fewer refractory connections. None of the motifs strongly delineate between biotic and abiotic system networks, though the parallel reaction motif m4 appears to show up slightly more often in biotic networks and refractory mx motifs appear to show up slightly less often in biotic networks.

### Chemical reaction networks span different regions of the chemical space

The networks selected for study are not necessarily isolated and individuated from one another: reactions found in one type of system (i.e., an oxidation process in an atmospheric reservoir) may have parallels in another (i.e., a respiration pathway in a biotic network). We quantified the chemical overlap between two networks as the number of molecules shared by two chemical networks divided by the smaller number of molecules of the two networks. This metric is 1 if one of the reaction networks contains all other chemical reaction network chemicals. Our results (**Figure 4a**) indicate two large endmembers in our dataset, biotic and atmospheric models, leaving the other chemical networks with just intermediate overlap values between these endmembers. Surprisingly, given its large size, the ORD chemical reaction network is more sparsely connected than we would have expected. Similarly, we computed the overlap of chemical reactions (**Figure 4b**), finding also that only the biotic and atmospheric networks are similar among themselves. Many of these networks are connected through molecular species, but they could contain completely different reactions depending on their domain.

**Figure 4:**
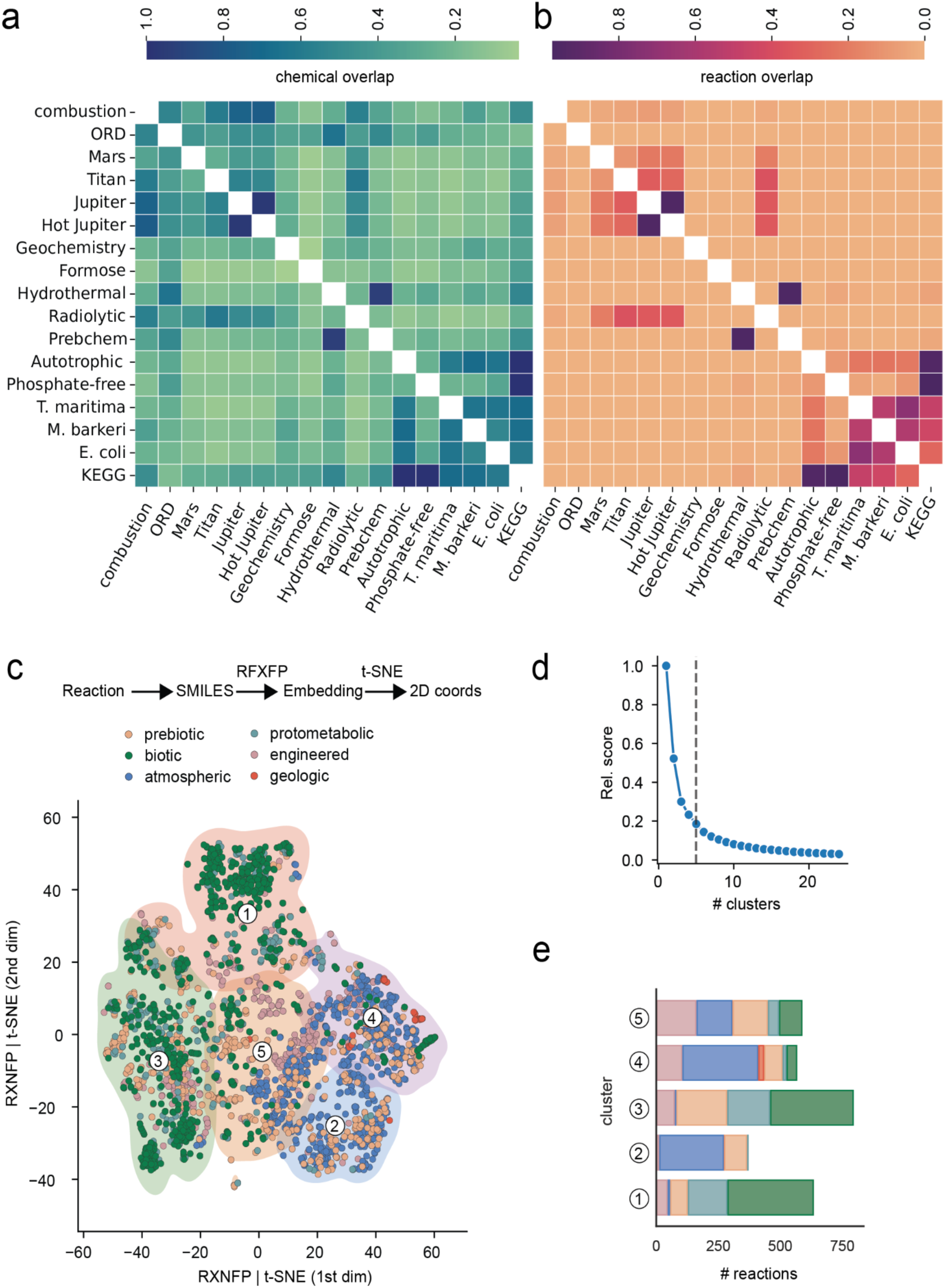
Chemical attributes across networks, excluding the *Translation* and *Cosmochemistry* networks due to their lack of computer-readable chemical reactions. **a)** Chemical overlap of each network pair, computed as the number of compounds in common divided by the number of compounds of the smallest network of each pair. **b)** hard-chemical overlap, computed as the number of identical reactions divided by the number of reactions of the smallest network of each pair. **c)** t-SNE representation of chemical reaction fingerprints obtained with RXNFP. Each marker represents a chemical reaction from one of the networks. **d)** Numbers represent clusters obtained with a K-means method, with approximated divisions given by the solid lines. The panel in the right shows examples of reactions close to the centroid of each cluster. **e)** A color-coded histogram chart of each cluster and the relative proportions of categorized reactions composing each cluster.

Although there is a near-infinite number of possible chemical reactions, most reactions can be reduced to a small number of similar mechanisms acting across and between different compounds (*e.g.* alkylation, esterifications, de-aminations, *etc.*). To understand if our networks use similar kinds of chemistry, despite not using the same exact reactions, we replaced the *hard similarity* of chemical reactions (*i.e.* transforming the same reactants into the same products) with a *softened similarity* metric based on the embedded distance between reaction-distributed representations^69^ .

We generated these embeddings of each chemical reaction using a transformer-based architecture trained in chemical reaction SMILES. As this technique is more computationally demanding, we employed it on a subset of ∼5,500 reactions chosen randomly but representing all the chemical networks. In **Figure 4c**, we plotted the embeddings after reducing their dimensionality through a t-SNE technique, which folds space to make similar objects lie close in the 2D plane. The result is a complex two-dimensional distribution with some apparent clusters of points. We computed the intra-cluster distance for different numbers of clusters (**Figure 4d**), and we found that a division of the distribution into five clusters provides the best choice, as this is the number of divisions at the elbow (the point in the distribution nearest the origin). In **Figure 4e**, we show the compositions of the different clusters by category of the networks from which the reactions were taken. Cluster 1 is composed mostly of biotic reactions. Clusters 2 and 4 are primarily composed of atmospheric/engineered reactions. Clusters 3 and 5 are composed of mixed sets of reactions. Prebiotic chemistry shows reactions in all 5 clusters, indicating that its chemistry likely borrows from or builds upon reactions found in different disciplinary domains of chemistry.

### Multidimensional analysis can impartially distinguish different network categories

The analyses above reveal that biotic chemical reaction networks tend to display certain features (heavy-tailed degree distributions, a low correlation between mass and degree, *etc.*), but those features are not entirely unique to biochemistry. For almost every attribute analyzed, one may find overlap with at least one other abiotic network type or category. For example, the ORD, radiolytic and cosmochemistry networks also have likely heterogeneous connection distributions. To assess all measured attributes impartially, we aggregated values from all attributes into a high-dimensional space and performed an Independent Component Analysis (ICA) with the full set of considered attributes displayed in **Table 5**. Only 15 networks for which all attributes could be measured could be included in the ICA analysis. This technique generates a representation of a multidimensional space in terms of just a small number of predefined axes that provide statistical independence (in contrast with *Principal Component Analysis*, which just rotates the axis to align them with the directions of maximum variance).

**Table 5.** Attributes employed in the Multi-Dimensional Independent Component Analysis.

| Attribute | Meaning |
| --- | --- |
| $\alpha$ | Exponent of the heavy-tailed degree distribution. |
| $\langle k \rangle$ | Average molecular degree / number of reactions involving each molecule in the network. |
| $r(\text{mass-k})$ | Pearson's correlation between molecule mass and degree |
| $m(1,2,3,4,5,x)$ | Frequency of motifs over the total of the network. |
| $\text{cluster}(1,2,3,4,5)$ | Clusters generated by t-SNE embedding |

Using two independent components, we were able to recover a clear separation between all networks based on their categorization, with a pronounced separation between the biotic and protometabolic networks (dark green and pale green) and all the other networks. The first component clearly separated atmospheric/geological networks and the biotic, protobiotic, and prebiotic networks.

The second component separated the prebiotic and the biotic networks, except for the radiolytic network, which lies closer to the atmospheric networks. An inspection of the component values of each of the vectors (**Figure 5b**) shows that the first component is mainly related with the degree-distribution parameter α, the average degree <K> and with the mx-motif frequency, so higher values along this axis represent more densely connected networks with higher degree-distribution exponents and a higher proportion of refractory motifs. The second component places more emphasis on the frequency of the shared product (m2) and parallel reaction (m4) micro-motifs, both of which were observed to occur with slightly greater frequency in biotic and protometabolic networks than in other abiotic networks.

**Figure 5.**
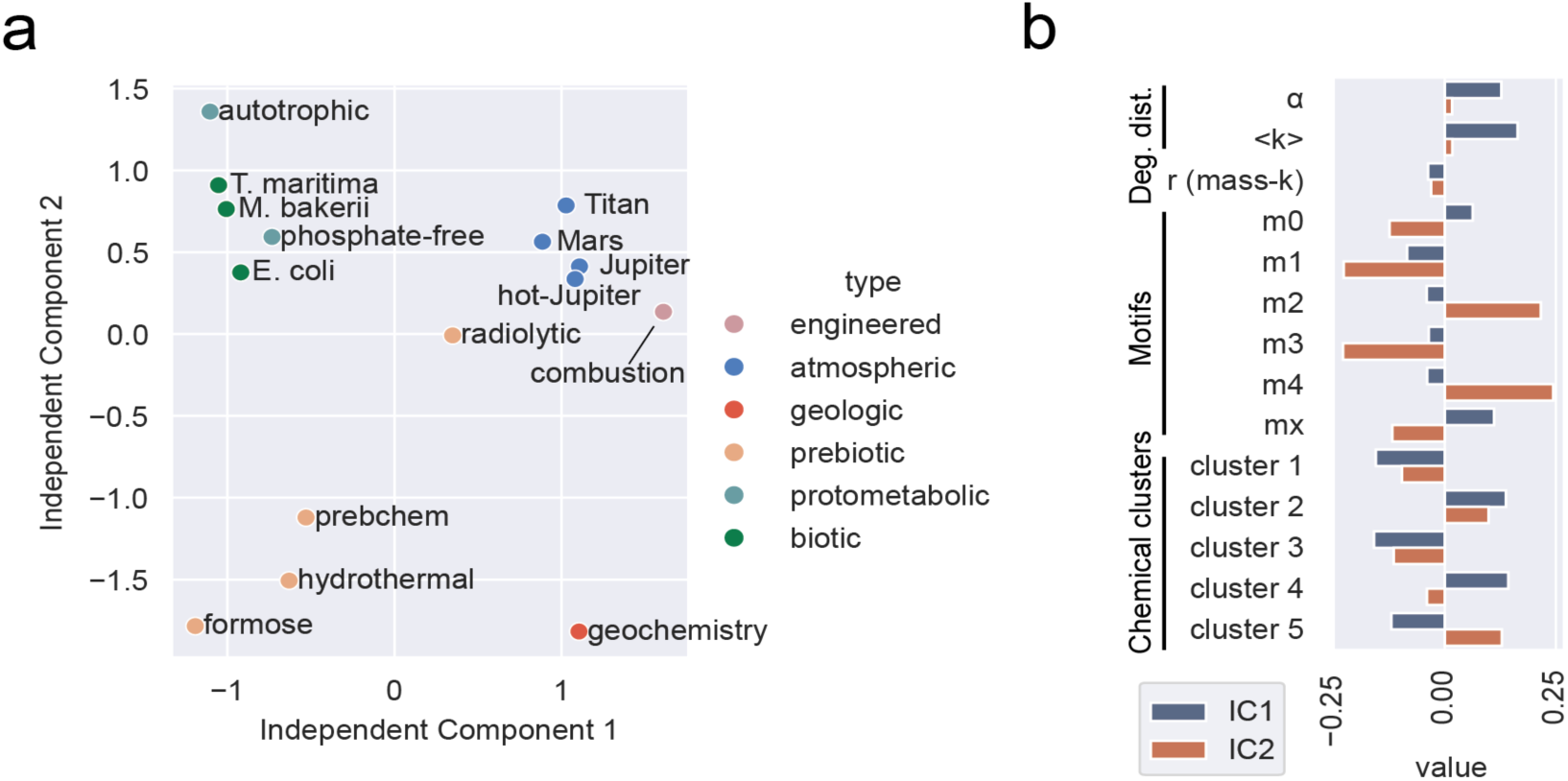
**a)** A two-dimensional Independent Component Analysis for all selected networks containing measurements of all network properties. Each point represents a network. The largest networks (KEGG, ORD, Cosmo-chemistry and translation) could not be characterized due to the incalculability of some of their attributes owing to computational limitations. **b)** A breakdown of the weights associated to each attribute on the two independent components. A positive value denotes a positive correlation to the component vector.

## Discussion

In this study, we compiled, annotated, and analyzed chemical reaction networks spanning a broad set of systems, from interstellar chemistry to biochemistry (see **Table 1**). Our goal was to identify features that impartially distinguish biotic from abiotic systems. The multidimensional analysis shown in Figure 5 demonstrates that a combination of both macroscale (network-level) and microscale (reaction-/compound-level) quantitative attributes contribute to this goal. Chemical reaction networks describing both biotic and abiotic networks in uniform ways were analyzed with the same tools, and their measured attributes aggregated, to obtain a two-dimensional representation. No single attribute by itself clearly delineates living systems, but linear combinations of some chemical reaction network attributes (*e.g.* degree distribution exponent, average degree, and specific micro-motifs) can separate biotic and abiotic systems. Note that the network category assignments that display as clustered colors in Figure 5a played no role in the ICA analysis-these clusters formed on the basis of measured CRN attributes alone.

Our study highlights significant differences between homogeneous and heterogeneous networks. Our measures of the degree distribution (**Fig 1**) provided a large difference between these two kinds of networks, with homogeneous networks being mostly atmospheric and heterogeneous networks being mostly biotic. We speculate three possible reasons for these differences: the fundamental differences between gas-phase and liquid-phase reactivity, the lack of evolutionary mechanisms in atmospheric chemistry, and the possibility that these differences are due to how these chemical reaction networks are built, which often depends on a thorough enumeration of all possible states and reactions of a molecule, even if their relevance is limited. The log-linear relationships between mass and degree (**Fig 2**) observed in homogeneous networks suggest that entropy-driven reactions and enumeration-based approaches influence this correlation. Specifically, in the gas phase, the increase of entropy may drive most reactions, leading to a large population of small, reactive molecules. Additionally, the enumeration-based approach used to generate atmospheric networks biases this distribution, as it becomes exponentially challenging to include the same description level with larger molecules. However, it is important to note that if only the enumeration-based approach were true, atmospheric models would not have been as useful since they would not allow the modeling of all relevant chemical spaces.

Our motif analysis (**Fig 3**) showed that most networks have a different distribution than expected on a random network, with parallel reactions and the absence of certain motifs indicating specific constraints and reaction patterns. For instance, many chemical networks may encode parallel reactions that reuse some of the reactants to produce similar products, such as biological reactions requiring cofactors like ATP or NADH. In atmospheric chemistry, such reactions could consume radiation or radicals. The absence of motif *m3* could be related to constraints that limit the recurrent reactivity of a given molecule with another one, and fully serial reactions might be more difficult to detect in certain settings like one-pot reactions. Motifs *mx* provide asymmetric relationships between molecules and reactions with a difficult rationale, yet they are still more abundant in all chemical reaction networks than other more symmetric motifs like *m1*, *m2*, or *m3*. Finally, our chemical similarity analysis showed that many of these networks are connected through molecular species, but they could contain completely different reactions depending on their domain. A softened version of this analysis indicates some affinity between the kinds of reactions in different networks, but with still clear divisions between atmospheric chemistry and biotic chemistry.

The development of this pipeline has direct implications for the fields of prebiotic chemistry and astrobiology. The CRNs gathered here were selected because they were readily available, and because they characterize as many different chemical systems as possible that are all candidates for having contributed to life’s origins or may serve as current targets for the search for life. From a categorical standpoint, for example, the atmospheres of very different bodies (Titan, Jupiter, Mars) all plot similarly to one another, and quite differently from either biotic or explicitly prebiotic organic synthesis networks. None of them seem obviously more or less predisposed to giving rise to life by comparison to one another, at least based on their CRN structures and content. In this sense, the networks and metrics gathered here quantify (in a limited way) how truly dissimilar biotic systems are from natural geochemical and atmospheric systems. By extension, these dissimilarities also quantify how far the prebiotic chemistry field may be from generating *de novo* chemical systems that resemble life. None of these CRNs, however, ought to be viewed as complete descriptions of their environments. Gathering more or different data would change the content of a given CRN and shift its plot point, perhaps sufficiently to trace a contiguous arc between these very disparate kinds of systems. The central plot location of the radiolytic CRN may indicate compatibility in knitting together reactions between and amongst these very different chemical systems. In any event, from a standpoint of chemosynthesis and CRN composition, it is clear that far more data are needed from prebiotic chemical systems research to significantly expand their connectivity to protometabolic and biotic systems.

The study also points to the development of new experimental studies based on topological properties alone, rather than composition. This view has been expressed in recent perspectives that have called for different approaches based on biology instead of the chemistry of life^70^ . Transitioning from studies designed to focus on the production of biomolecules to the emergence of behavioral attributes afforded by proto-cells, proto-metabolisms or proto-genomes requires significant conceptual steps ^71^ . As just one example of this challenge, the biomolecules that we know could result from the evolution of metabolism^72^ , posing questions about the timing and chronology of current biomolecules.

Our study dives into this problem from a different perspective, considering not the specific molecules or reactions that should make life possible but rather the properties of the resulting networks. For instance, our study indicates that power-law-shaped degree distributions with an exponent around 2.0 or a low micro-motif density for mixed motifs (mx, in **Figure 3**) are attributes associated with networks describing living systems. Previous research has shown how processes such as radiolysis^66,73^ , environmental factors^74^ , or wet-dry cycles ^24^ could drive the topological features of chemical reaction networks without involving specific catalysts. Controlling the interaction patterns between reactions could be a more challenging target, but recent implementations of complex molecular networks in DNA strand displacement reactions ^75^ show a promising platform for expanding topological studies of organic CRNs.

All models are simpler than reality, and by extension all real-world systems are more complicated than the networks that describe them. In this study, the assembled CRNs represent one possible model that captures many of the observed and inferred chemical facets of these different environments. This simplification means that we have neglected stoichiometry (required to find autocatalytic cycles), thermodynamics (required to find reaction directionality), and kinetics (required to understand the time evolution of the system). These limitations emerge from the challenges of obtaining these data. For instance, most condensed-phase chemical reactions are discovered in complex chemical mixtures, so the stoichiometry of the final products is not apparent. Finding stoichiometries could have enabled us to perform computational predictions based on first-principles calculations —although automating accurate unsupervised and *ab initio* calculations can also be challenging in its own right. A promising alternative and possible way of advancing these techniques may be found in the usage of machine-learning models based on fuzzy representations of the chemical reactions without requiring their full stoichiometric description^76–78^ .

Another area for improvement of this study is its small molecule focus, which might disregard the role of larger molecules, mesoscopic molecular phases, and macromolecules in shaping the chemical reaction networks of life. Indeed, previous research has indicated the possibility of life requiring mass-discontinuities through its components ^79^ ; a phenomenon that we cannot describe in this work in detail as we lack the information about macromolecule or grain formation in each of the systems. Future theories and experiments could highlight the role of system dynamics in forming macro-aggregates and macromolecules, together with the size distribution of those chemical entities. However, whether those approaches could be combined with our small-molecule-focused studies remains an open question.

## Conclusions

The problems facing the study of life’s origins are in parallel to those faced by fundamental problems in biology. Both efforts are undertaken in an attempt to observe and understand how genuine multi-level complexity can be implemented and preserved over time in a chemistry-based, self-organizing system. Prebiotic chemistry, though, is lacking in any single concrete model system that links entirely abiotic chemistry to entirely biotic chemistry, by which the development of new experimental designs and hypotheses may be guided. The sheer number of behavioral ways in which living systems differ from non-living systems, and a poor understanding of the relative chronology and co-dependence of structural features that enable these behaviors to occur, represents a formidable and enduring challenge.

The focus explored here, with its acknowledged limitations, provides insights into the basic features that a biology-based chemical reaction network ought to display that would distinguish it from an array of abiotic chemical reaction networks. As such, measurement of these features may be employed as a basic guide for assessing whether a novel candidate system may be capable of giving rise to life, and also for impartially assessing whether an existing prebiotic candidate system is comparable to biotic systems than to other candidate systems under consideration for study. The array of quantitative features used in this kind of assessment can and likely will grow as the field matures and as observational techniques improve. Perhaps the most surprising finding is that detecting and distinguishing life as a network of chemical reactions does not obviously require the development of entirely new metrics, despite the widely acknowledged etymological difficulty faced by those attempting to precisely define what life is.

